# Updated population viability analysis, population trends and PBRs for Hector’s and Maui dolphin

**DOI:** 10.1101/2020.03.25.008839

**Authors:** Elisabeth Slooten, Stephen Michael Dawson

## Abstract

Updated population viability analyses, incorporating the latest abundance and bycatch data indicate that:

- The estimated population decline for Maui dolphin is 2% per year
- There is a 68% probability that the population is continuing to decline
- After 30 years (6 more surveys) statistical power of detecting this rate of decline would be < 15%
- Only very large declines (37%) and recovery (45%) would be detectable with 80% statistical power after 30 years (6 more surveys)
- The level of conservation threat for Hector’s dolphin remains high despite a recent, larger population estimate off the east coast of the South Island
- Increased overlap between dolphins and fisheries, due to more extensive offshore distribution of dolphins off the South Island east coast, more than offsets the apparently higher population size
- The Hector’s dolphin population has declined 70% over the last 3 generations, exceeding the 50% threshold for Endangered
- Population declines are predicted to continue under current protection levels
- The results of this research are consistent with:
  ◯ NOAA proposal to list Hector’s and Maui dolphins under the US Endangered Species Act
  ◯ IWC recommendation to ban gillnets and trawling throughout Maui dolphin habitat
  ◯ IUCN recommendation to ban gillnets and trawling throughout Hector’s and Maui dolphin habitat

## INTRODUCTION

This paper provides updated analyses of abundance and catch data for Maui and Hector’s dolphin, including a Population Viability Analysis (PVA) and estimates of Potential Biological Removals (PBR). This work was carried out in response to a request from the United States National Oceanographic and Atmospheric Agency (NOAA) to support the process of considering the listing of Hector’s dolphin as threatened and Maui dolphin as endangered under the US Endangered Species Act. NOAA requested additional information from the scientific community, in particular new or updated data on threats and population viability analyses for Hector’s and Maui dolphins.

We use the most recent estimates of abundance and fisheries mortality to update population viability analyses for Hector’s and Maui dolphins. The most recent published population viability analyses (Slooten 2007; Slooten and Dawson 2010) used population estimates from the first series of line-transect surveys, conducted during 1997-2001. These surveys ranged 4 nautical miles offshore and resulted in an estimate of 7,270 Hector’s dolphins (95% = 5,303 – 9,966) and 111 Maui’s dolphins (95% CI = 48-252, CV=44%, Slooten et al., 2004). A second series of line-transect abundance surveys, ranging to 20 nautical miles offshore, resulted in an estimate of 14,849 Hector’s dolphins (95% CI = 11,923 – 18,492). A more recent estimate for Maui dolphins is also available, of 63 dolphins one year and older (CV 9%) from genetic mark-recapture methods (Baker et al. 2016). We have used the same methods as previous PVAs (Slooten 2007; Slooten and Dawson 2010) in order to explore how conclusions about risk and population viability are affected by the new data.

Population estimates from the most recent line-transect surveys, conducted during 2010-2015, were very similar to estimates from the previous surveys in all areas except the east coast of the South Island. A series of three summer and three winter surveys off Banks Peninsula showed that Hector’s dolphins range offshore to the 100 m depth contour (Rayment et al. 2010). The offshore distance of this contour explains differences in distribution of Hector’s dolphins among areas. Off parts of the South Island east coast where the bottom shelves very gradually (e.g. South Canterbury), Hector’s dolphins range out to approximately 20 nautical miles (n.mi) offshore.

Issues with field protocols and analysis methods appear to have contributed to the relatively high population estimates. Some of these problems appear to be generic, such as occasional extremely high estimates or “blow outs” associated with the use of Mark-Recapture Distance Sampling (IWC 2016). Other problems are specific to this survey (e.g. incomplete overlap between observers). The Scientific Committee of the International Whaling Commission has encouraged further work in response to the recommendations it its report on how to improve the field protocols and analysis methods (IWC 2016). In this PVA we take the new abundance estimates at face value. This should not be taken as an endorsement of them.

The level of fisheries mortality in gillnet and in particular trawl fisheries is the model input with the highest level of uncertainty. This is due to very low and sporadic observer coverage in New Zealand’s inshore fisheries. We briefly review the available data on fisheries mortality. The first observer programme, carried out in 1997-98 (Baird and Bradford 2000), remains the most scientifically robust. New observer data are sparse, but consistent with earlier estimates.

For these analyses, we have taken the most recent estimates of dolphin population size and bycatch at face value. We look forward to re-running the analyses described in this paper as more scientifically robust estimates become available.

## MATERIALS AND METHODS

Current protection measures (also see Figure 1):

**Figure 1.**
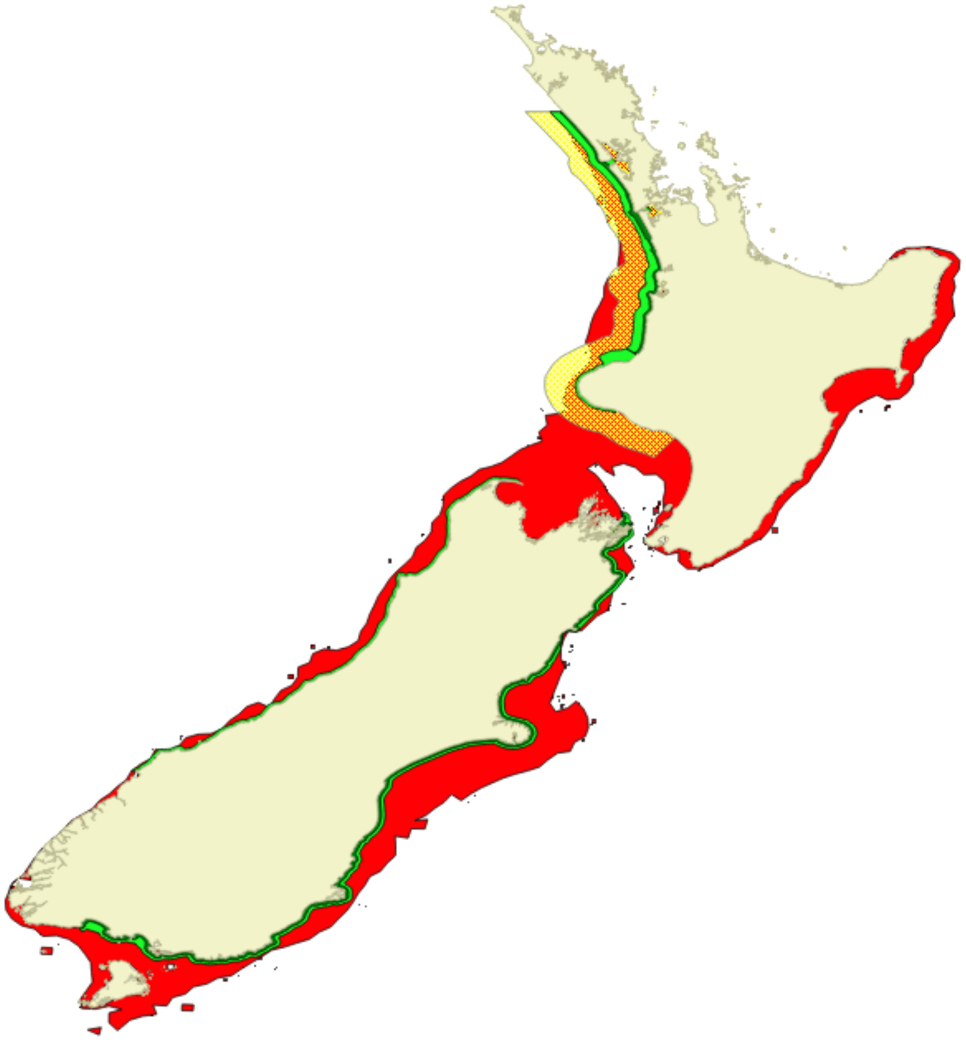
Map of Hector’s dolphin (South Island) and Maui dolphin (North Island) range out to the100m depth contour (red area). Protection from gillnet fisheries (light green) and both gillnet and trawl fisheries (dark green) is shown as well as the level of protection recommended by the International Whaling Commission (gillnet and trawl ban in yellow area). The IUCN recommend protecting the entire habitat of Hector’s and Maui dolphins (red area) from gillnet and trawl fisheries.

### WCNI

West Coast North Island: Maunganui Bluff to Whanganui

- Commercial gillnetting is banned off open coasts to 7 n.mi offshore from Maunganui Bluff to Pariokariwa Point and to 2 n.mi from Pariokariwa Point to Hawera. No protection from gillnets from Hawera to Whanganui.
- Commercial and amateur gillnetting is banned in the entrances of the major harbours, but is legal inside harbours.
- Trawling is banned within 2 n.mi offshore from Maunganui Bluff to Pariokariwa Point, and to 4 n.mi between Manukau Harbour and Port Waikato. No protection from trawling from Pariokariwa Point to Whanganui.

### ECSI and SCSI

East Coast and South Coast of the South Island:

Cape Jackson in the Marlborough Sounds to Sandhill Point east of Fiordland

- Commercial gillnetting is banned from the coastline to 4 n.mi offshore, except at Kaikoura, where it is banned to 1 n.mi offshore, and in Te Waewae Bay, where it is banned to approximately 9 n.mi offshore.
- Amateur gillnetting remains allowed in some harbours, estuaries and inlets. For example, amateur gillnetting for flounder is permitted between 1 April and 30 September in the upper reaches of four harbours on Banks Peninsula, and in shallow waters of Queen Charlotte Sound.
- Trawling within 2 n.mi of shore is allowed, using nets with < 1.5 m headline height.

### WCSI

West Coast South Island: Cape Farewell to Awarua Point

- Commercial gillnetting is banned within 2 n.mi offshore between 1 December and 28 February.
- No protection from trawling.

### NCSI

North Coast South Island: Cape Farewell to Cape Jackson

- No protection from gillnet or trawl fisheries.

New Zealand’s national progress reports to the Scientific Committee of the IWC provide reported bycatch for the period 1985-2015 (IWC 2016). Additional data are available from a Department of Conservation database (DOC 2016), the scientific literature and official government reports (Abraham et al. 2010, 2016; Abraham and Thompson 2011; Baird and Bradford 2000; Berkenbusch et al. 2013; Currey et al. 2012; CSP 2015; Davies et al. 2008; Dawson 1991; MPI 2013, 2016; Ramm 2010, 2012; Rowe 2009, 2010; Slooten 2013, 2014; Slooten and Davies 2011; Starr and Langley 2000; Thompson et al. 2013; Abraham et al. 2016). Bycatch is known to occur in trawl fisheries, but observer coverage is currently too low to allow robust estimation of the catch rate. Likewise, there are no recent estimates bycatch in recreational gillnet fisheries. Gillnets can be legally used by recreational or amateur fishers for most (or all) of the year in South Island harbours and year-round in the harbours of WCNI.

To assess the impact of mortality in the commercial gillnet fishery, we carried out a population viability analysis, following the same methods as Martien *et al*. (1999), Burkhart and Slooten (2003), Slooten (2007) and Slooten and Dawson (2010). We used a standard surplus production model:

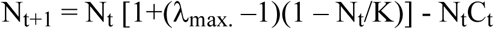

Where N_t_ = population size at time t, λ_max._ = maximum annual population growth rate, K = population size in 1975 and C_t_ = the catch rate in year t, estimated on the basis of fishing effort in year t and data from observer programmes (as outlined below).

For each run of the model, a maximum population growth rate was selected from a uniform distribution between 1.018 and 1.049 to reflect uncertainty in the estimation of this parameter. The highest and lowest population growth rates span the range estimated by Slooten and Lad (1991) and also used by Martien *et al*. (1999) and represent a highly optimistic parameterisation. The smallest value (1.018) is the estimated λ_max._ for Hector’s dolphin, based on field data (Slooten and Lad, 1991). The highest value (1.049) is an estimate of the maximum possible λ_max._ for the species, based on survival curve for humans and the most optimistic estimate of reproductive rate for Hector’s dolphin (Slooten and Lad, 1991). Using a uniform distribution between 1.018 and 1.049 means that values at the upper, highly optimistic end are equally likely to be used as values at the lower, more realistic end. Hector’s dolphin, like most other small cetaceans, have low potential population growth rates (e.g. Perrin and Reilly, 1984: Reilly and Barlow, 1986). Another optimistic aspect of the model is that it assumes the population growth rate is highest in very small populations. It does not take into account the potential for Allee effects – decompensation at very small population sizes, e.g. due to inbreeding, difficulty of finding mates, reduced effectiveness in feeding or predator defense.

We used the most recent estimates of population size (MacKenzie and Clement 2014, 2016, Baker et al. 2016). For each of 5000 back calculations, values for N and λ_max._ were randomly selected from their respective distributions and population size in 1975 was estimated by back calculation. This is a common approach used in fisheries and marine mammal assessments (e.g. Smith and Polachek 1979, Barlow and Hanan 1995, Martien *et al*. 1999). Forward projections used these 5000 sets of values, each set maintaining the relationship between K, N and λ_max._ for each forward projection of the model. For more detailed information on the methods used for the PVA please see Slooten and Dawson (2010).

The IUCN has listed Hector’s dolphin as Endangered based on the basis of an estimated > 50% population decline over the last three generations (39 years, IUCN 2016). Therefore, we compared current (2013-2015) abundance estimates with estimated abundance in 1975. Gillnet fisheries in New Zealand waters expanded rapidly in the late 1960s and early 1970s. Therefore we are confident that original population size was higher than 1975 levels.

We also update PBR estimates for Hector’s and Maui dolphins. The PBR method (Wade 1998) aims to ensure that human-caused mortality is below levels that could lead to population depletion. It is based on a logistic model with maximum net productivity level (MNPL) at 0.5K. MNPL is the population size that results in the maximum number of individuals being added to the population per year. At MNPL one would expect the population growth rate to be approximately 0.5 R_max_ (Slooten and Dawson 2008). As the goal is to ensure populations stay at or above MNPL (i.e. above 0.5 K), the population growth rate used in the PBR calculation is 0.5 R_max_. For marine mammals MNPL is thought to be between 0.5 K and 0.85 K (Taylor and de Master 1993).

The method explicitly takes into account uncertainty and potential biases in the available information. PBRs were calculated using using the following equation:

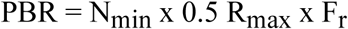

Where N_min_ = 20^th^ percentile of the population size estimate, R_max_ = Maximum annual population growth rate and F_r_ = Recovery factor.

In the design of the PBR method, a range of mortality limits were evaluated based on whether at least 95% of the simulated populations met two criteria: (1) populations starting at MNPL stayed at or above MNPL after 20 years, and (2) populations starting at 0.3 of K recovered to at least MNPL after 100 yr (Wade 1998). Simulations indicated that using the 20^th^ percentile of the population size estimate met those criteria (Wade 1998). N_min_ estimates for Hector’s dolphin populations from the most recent population surveys (MacKenzie and Clement 2016, Baker et al. 2016) were used to calculate PBR. When data on productivity are available it is recommended that they be used (Wade 1998). R_max_ has been estimated at 0.018 based on marine mammal survival rate curves and the most optimistic reproductive parameters (age at first reproduction and calving interval) estimated in the field (Slooten and Lad 1991). Here we calculate PBRs using the R_max_ estimate for Hector’s dolphin (0.018). PBRs based on the default value for cetaceans in general (0.04) are also included for comparison.

Simulations that included plausible levels of bias in the available information indicated that F_r_ needs to be ≤ 0.5 to meet performance criteria (1) and (2) above (Wade 1998). Taylor and Wade (2000) showed that using N (rather than N_min_) and F_r_ 1.0 resulted in many of the simulated populations being depleted below 0.5 K. They also carried out robustness trials, similar to those used by the Scientific Committee of the International Whaling Commission in testing its Revised Management Procedure (Donovan 1989). Two plausible flaws in the data or assumptions were explored, based on biases (e.g. in abundance and mortality estimates) similar to those observed in the management of marine mammal bycatch (Taylor and Wade 2000). For endangered species an F_r_ of 0.1 is recommended (Wade and Angliss 1996; Wade 1998). For example N_min_ for North Atlantic right whale is 476 individuals, F_r_ is 0.1 because it is Endangered and the PBR is 1 individual per year (NMFS 2015). With a PBR this low “any mortality or serious injury for this stock can be considered significant” and the management goal is to reduce fisheries mortality to as close to zero as practicable (Fisheries and Oceans Canada 2014, NMFS 2004, 2015).

For Maui dolphin (WCNI population) we estimated population trends, the probability of continued population decline and the statistical power of being able to detect population trends from continued population surveys. The Maui dolphin population has been declining at c. 3% per year since 1985, from 140 individuals in 1985 (95% CI 46-280; Dawson and Slooten 1988) to 55 individuals in 2011 (95% CI 48-69; Hamner et al. 2012). The last three population estimates were 69 (95% CI 52-100; Baker et al. 2012), 55 (95% CI 48-69; Hamner et al. 2012) and 63 (95% CI 57-75; Baker et al. 2016). This raises the question as to whether the last estimate is consistent with a continuing decline of 3% per year and how many more surveys would be required to provide a conclusive answer to this question.

To compare population estimates statistically, we have used both frequentist and Bayesian approaches. In calculating trends we have included all abundance estimates with the exception of Russell (1999). That survey did not develop its own correction factors for offshore distribution or fraction missed, but assumed those values from Dawson and Slooten (1988). The remaining 6 surveys used a variety of methods. We have followed Wade et al (2012) in treating all these at face value and fitting a linear regression to the natural log of the abundance estimates. To investigate evidence for a difference between the two most recent estimates, we also calculated a frequentist 95% confidence interval of the difference. We also present a Bayesian linear regression of the last three estimates, carried out in OpenBugs (Lunn et al. 2009). The posterior distribution from 20,0000 iterations (4 chains, 5000 iteration burn-in period) was used to estimate the probability of population increase or decrease.

A power analysis (Taylor and Gerrodette 1988) was used to estimate the statistical power of being able to detect future population declines or recovery. The power analysis used the function Powertrend in the R package Fishmethods (Nelson 2014). This package conducts power analysis for detecting trends following the methods of Gerrodette (1987, 1991, 1993) and Gerrodette and Brandon (1993). Powertrend allows the user to select a linear or exponential trend. We used the exponential option, to estimate the statistical power of detecting different rates of population decline and recovery, depending on the number of surveys conducted. We estimated the statistical power of being able to detect a continued population decline of 3% (the rate of decline estimated by Wade et al. 2012) and a population recovery of 1.8% per year (the maximum realistic rate of recovery estimated by Slooten and Lad 1991). Our power analyses estimated the probability of continued population surveys being able to detect these rates of population change. The analyses use the most recent population estimate of 63 Maui dolphins (1 year and older) in 2015-2016 and its CV (0.09; the smallest CV in the series of population estimates to date) as the starting point. The most recent population surveys for Maui dolphins have been 5 years apart and the government agencies funding the surveys have indicated that this will be the timeframe for future surveys. We estimated the statistical power of being able to detect specific rates of population change, as well as the rate of decline and recovery that would be detectable with 80% statistical power.

## RESULTS

### Bycatch

A total of 139 NZ dolphin entanglements were reported in New Zealand’s progress reports to the IWC for the period 1985-2015. On average, 4.5 NZ dolphin entanglements were reported per year, with higher numbers during periods when interviews with fishers were conducted (1985-1988) or observer programmes (e.g. 1998) were in place. In years when separate statistics were provided, the number of reported Maui dolphin entanglements was 2, 3, 2 and 1 in 2000, 2001, 2002 and 2012 respectively. Three trawl entanglements of Hector’s dolphin were reported in 2006. The New Zealand Department of Conservation (DOC) maintains a Hector’s and Maui dolphin incident database (DOC 2016). These data are presented by year in Table 2, for the same time period as the data in Table 1.

**Table 1:**
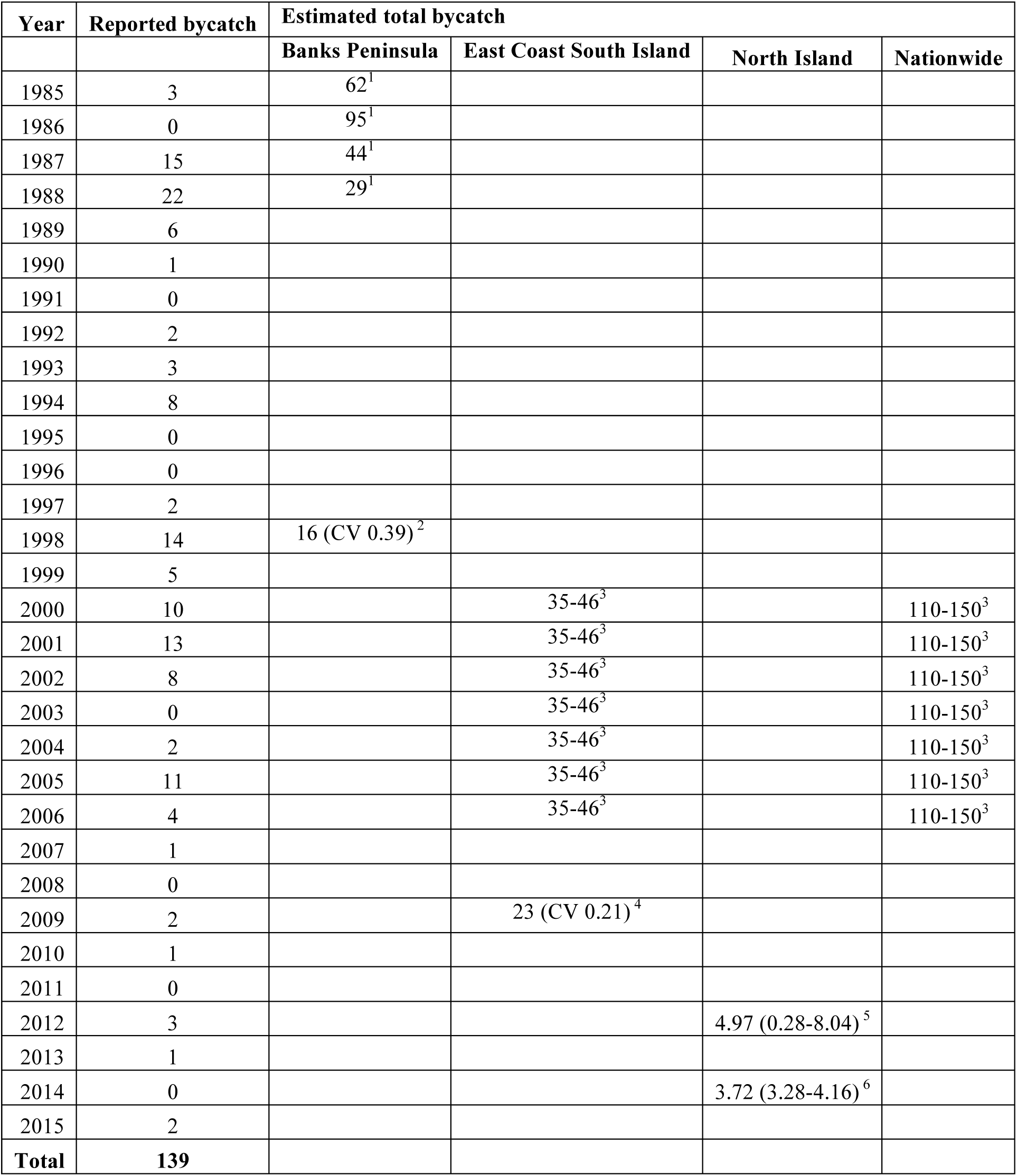
Number of reported Hector’s and Maui dolphin entanglements in fishing gear in New Zealand’s national progress reports to the IWC Scientific Committee. In most cases, the year in column one is the calendar year. Data sources are: ^1^ interviews with fishers (Dawson 1991); ^2^ Estimate from observer programme; ^3^ estimate based on 1997-1998 observer programme strike rate and 2000-2006 fishing effort (Davies et al. 2008); ^4^ 2008-2009 observer programme (Slooten and Davies 2011); ^5^ Expert Panel (Currey et al. 2012); ^6^ Expert Panel estimate modified for increased area protected (Slooten 2014).

**Table 2.** Data from the DOC database. The “other” causes of death column includes boat strikes, gunshot, live strandings and euthanased dolphins.

| Year | Gillnet | Trawl | Craypot | Possible bycatch | Other | Natural | Unknown |
| --- | --- | --- | --- | --- | --- | --- | --- |
| 1985 | 6 |  |  | 1 |  |  | 6 |
| 1986 | 12 | 1 |  | 1 |  |  | 2 |
| 1987 | 11 | 1 |  | 6 |  |  | 4 |
| 1988 | 8 | 4 |  | 3 |  |  | 14 |
| 1989 | 6 |  | 1 | 2 |  |  | 8 |
| 1990 | 1 |  |  | 2 | 1 |  | 7 |
| 1991 | 1 |  |  |  |  |  | 5 |
| 1992 | 2 |  |  | 5 |  |  | 11 |
| 1993 | 2 |  |  | 2 |  |  | 7 |
| 1994 | 4 |  |  | 5 |  |  | 7 |
| 1995 | 1 |  |  | 4 |  |  | 10 |
| 1996 | 4 |  |  | 2 |  |  | 14 |
| 1997 | 1 | 1 | 1 | 2 |  |  | 7 |
| 1998 | 10 | 1 |  | 6 | 1 |  | 6 |
| 1999 |  | 1 |  | 8 |  |  | 4 |
| 2000 | 2 |  |  | 7 |  |  | 8 |
| 2001 | 4 |  |  | 6 |  |  | 13 |
| 2002 | 8 |  |  |  |  | 4 | 12 |
| 2003 | 2 |  |  | 2 |  | 1 | 9 |
| 2004 | 1 |  | 1 | 1 |  | 3 | 9 |
| 2005 | 9 |  |  | 7 | 1 | 4 | 2 |
| 2006 | 1 | 3 |  |  |  | 3 | 7 |
| 2007 | 1 |  |  | 3 |  | 5 | 15 |
| 2008 |  |  |  | 2 |  | 3 | 12 |
| 2009 | 1 |  |  |  |  | 2 | 9 |
| 2010 | 2 |  |  |  |  | 3 | 5 |
| 2011 |  |  |  |  |  | 3 | 4 |
| 2012 | 6 |  |  |  |  | 8 | 12 |
| 2013 |  |  |  |  |  | 7 | 5 |
| 2014 | 1 |  |  |  | 5 | 3 | 2 |
| 2015 | 3 |  |  |  | 1 |  | 5 |
| **Totals** | **110** | **12** | **3** | **77** | **9** | **49** | **241** |

Of the 501 NZ dolphins in the DOC database, 290 were attributed a natural or unknown cause of death and 211 individuals were attributed to bycatch, possible bycatch or other causes (including boat strikes, gunshot, etc). Of these 211 dolphins, 52% were caught in gillnets and 96% were attributed bycatch or possible bycatch as the cause of death. There were 12 reported dolphin captures in trawl nets between 1985 and 2015. In this database, an additional 7 Hector’s and Maui dolphins were reported caught in trawls during the 1970s, including one incident with 3 dolphins caught in a single trawl and one incident with 4 dolphins in a single trawl. The high proportion of multiple captures complicates estimation of dolphin mortality in the trawl fishery.

Most of the discrepancies between the datasets relate to whether they include entanglements reported by observers, fishermen and/or dolphins found beachcast. For example, two Hector’s dolphins found beachcast in a gillnet (Figure 2) were included in the DOC database, but not in reports to the IWC. A Hector’s dolphin found beachcast in Akaroa Harbour is included in the DOC database as “probable bycatch” on the basis of clear net marks on the dolphin’s snout (Figure 2), but was not included in New Zealand’s report to the IWC. Video camera monitoring (MPI 2013) showed a fisherman catching two Hector’s dolphins, leaving the net in the water for an hour until the second dolphin dropped out of the net and reporting only the dolphin brought on board in view of the video camera (MPI 2013).

**Figure 2.**
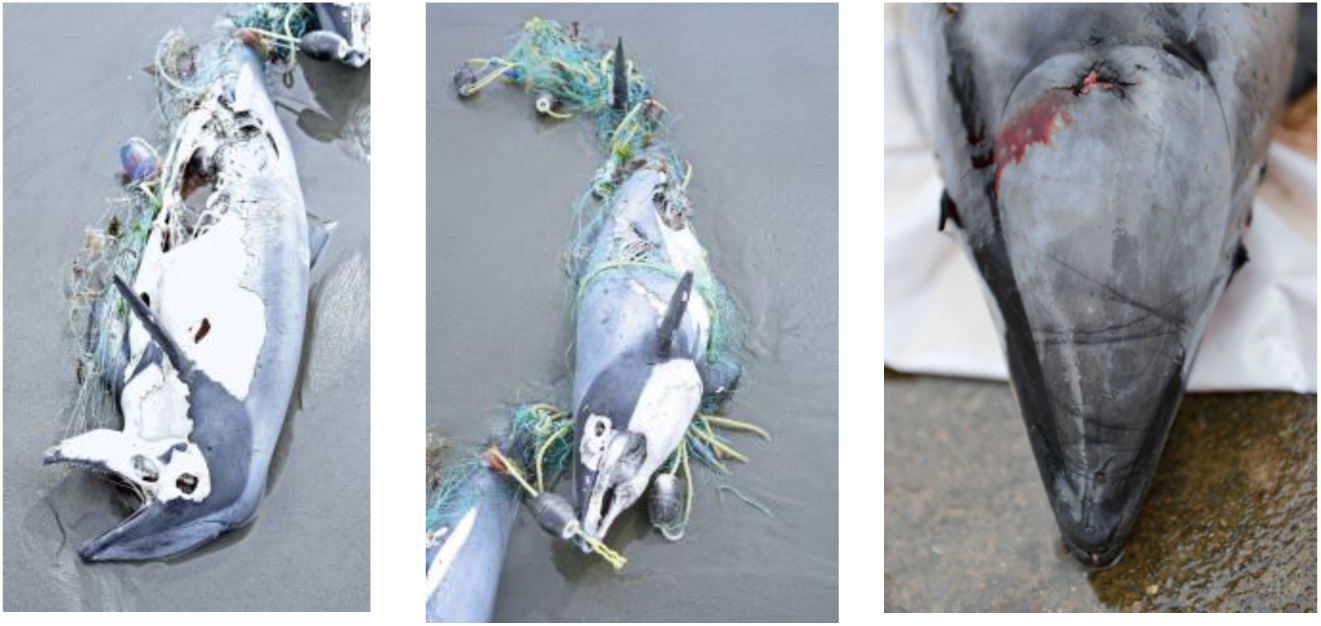
**Left and middle:** Two Hector’s dolphins found entangled in a gillnet, in 2012, included in the DOC database, but not reported in New Zealand’s National Progress Report to the IWC. **Right:** Net marks on Hector’s dolphin found beachcast in 2015, included as “probable” bycatch in DOC database but not reported to IWC.

Overall, the number of bycatch events reported to the IWC (139) is similar to the number of dolphins listed as “known bycatch” in the DOC database (125), but substantially lower than the number scored as known, probable and possible bycatch (211). Diagnoses appear to have become more conservative over time, with a decreasing proportion of dolphins diagnosed as “bycatch” and increased numbers attributed to “unknown” cause of death and “natural causes” (Figure 3). Some of the dolphins listed as “natural causes” were too decomposed to draw robust conclusions about cause of death. For example, the necropsy results for one of these dolphins states that “the state of decay of this body makes it difficult to determine the cause of death” the cause of death was reported as “natural causes”.

**Figure 3.**
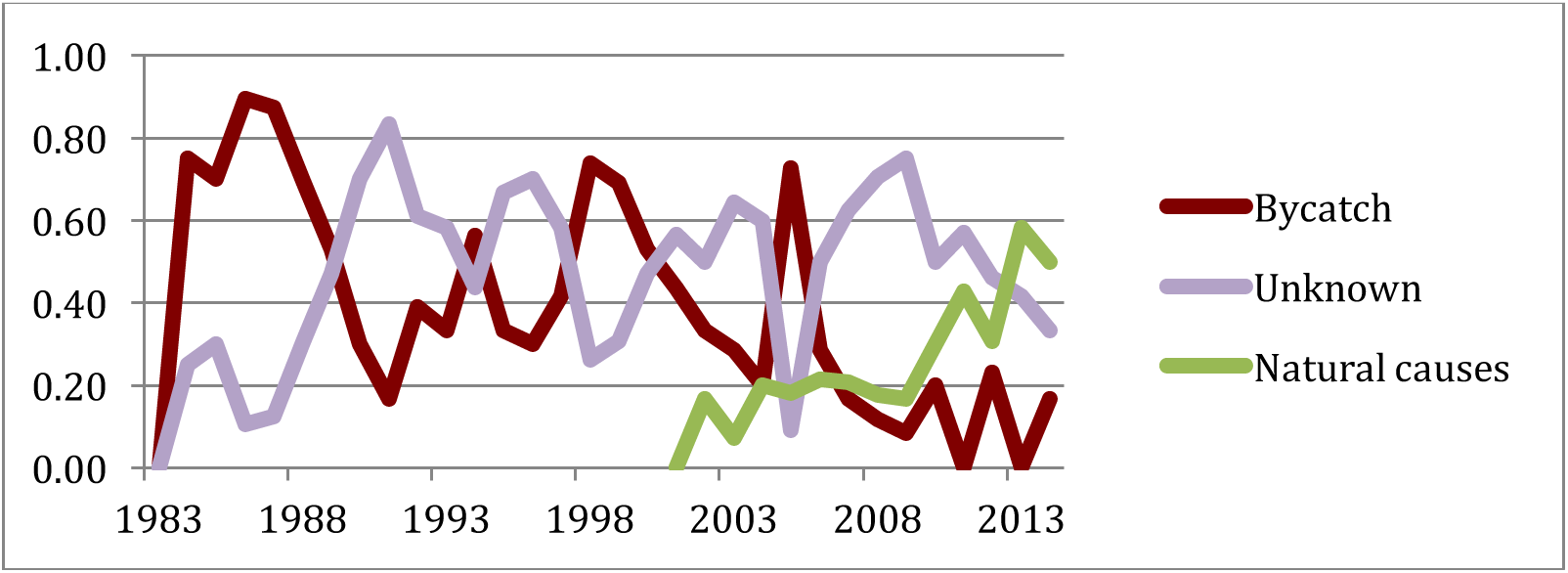
Summary of the data in the DOC database, showing the proportions of individuals with bycatch, natural or unknown cause of death.

Government reports typically provide numbers of “reported” dolphin deaths rather than estimates of the total number of dolphins caught. Estimates of bycatch based on dolphins found stranded are typically substantial underestimates. Only a small proportion of dolphins caught in fisheries subsequently strand on beaches, and are then found and reported to government authorities. In addition, not all dolphins killed in fishing gear show clear signs of the cause of death. Therefore some of the dolphins listed as having an “unknown” or “natural” cause of death are likely to have been caught in fishing gear. We have autopsied 119 NZ dolphins of which 83 were caught in gillnets, 5 in trawl nets and 31 were found beachcast with no obvious cause of death. Dolphins caught in trawl nets typically showed no signs of entanglement, except for contusions. On average, about 50% of the NZ dolphins brought in by fishermen or observers, and known to have been caught in a gillnet, showed clear evidence of entanglement.

More reliable data are available from observer programmes. However, the level of observer coverage has typically been very low. Figure 4 provides an indication of the level of observer coverage and the fact that bycatch is essentially undetectable at very low levels of observer coverage. The ECSI area has had relatively high levels of observer coverage, compared to other parts of the country, but still well below 10% in most years. A relatively higher proportion of dolphin catches have been reported as “released alive” in recent years (Figure 5), following a similar trend to the “natural causes” determinations for the dataset of beachcast dolphins (Figure 3). In summary, the official bycatch data show several biases,which act to underestimate the level of fisheries mortality. These problems underscore the need for a robust observer programme.

**Figure 4.**
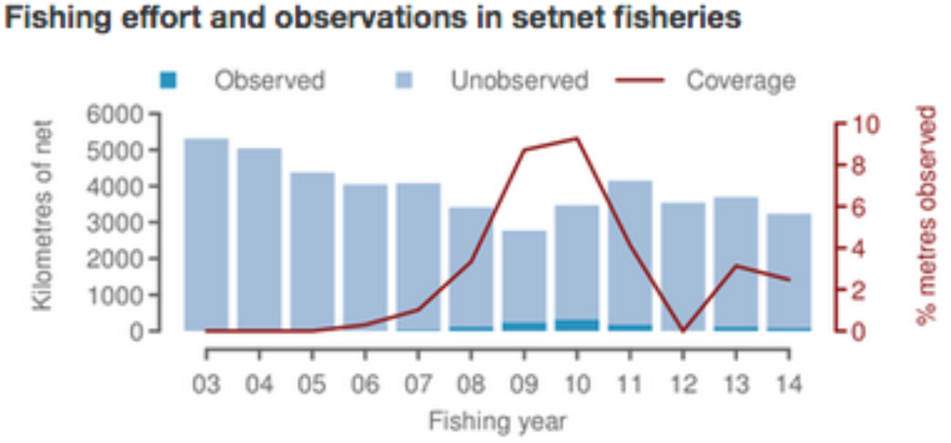
Level of observer coverage for ECSI from 2003-2014, downloaded from Dragonfly Consulting (Abraham et al. 2016).

**Figure 5.**
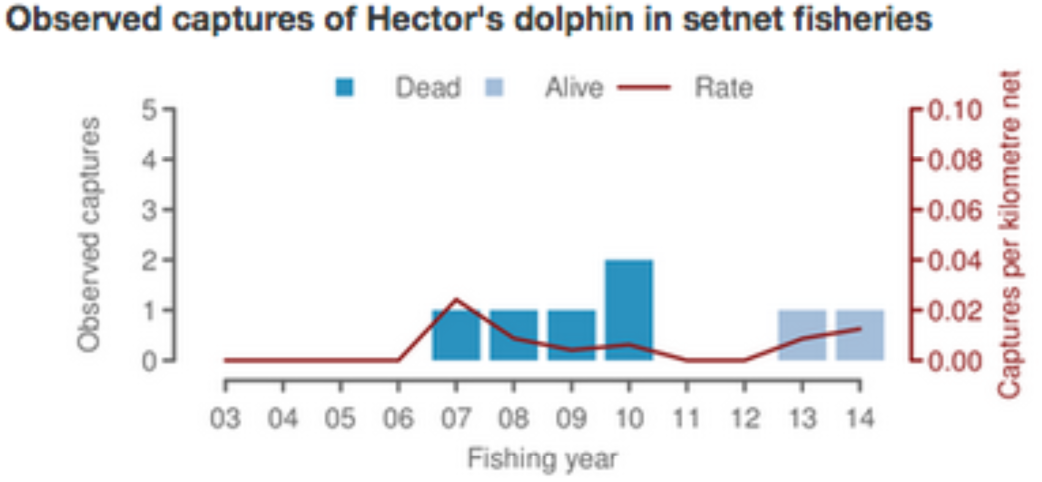
Summary of bycatch data for ECSI from 2003-2014, downloaded from Dragonfly Consulting (Abraham et al. 2016).

In 2000, the National Institute of Water and Atmospheric Research (NIWA) published the first estimate of Hector’s and Maui dolphin bycatch for gillnet fisheries around New Zealand (Davies et al. 2008) based on catch rates from an observer programme around Banks Peninsula (Baird and Bradford 2000). NIWA and fishing industry scientists estimated nationwide bycatch in gillnet fisheries at 110-150 Hector’s and Maui dolphins per year during the period 2000-2006 (Davies et al. 2008). Of these catches, 5-6% were included in New Zealand’s National Progress reports to the IWC and 1-15% were included in the DOC database. Davies et al. (2008) estimated ECSI bycatch at 35-46 Hector’s dolphins per year. Slooten and Davies (2011) estimated that protection implemented in 2008 decreased bycatch off the ECSI to 23 Hector’s dolphins annually. ECSI is the only area for which data from before and after 2008 are available, allowing such a comparison.

The highest level of observer coverage achieved to date was 22.6% in the observer programme conducted during the 1997-1998 fishing season (Baird and Bradford 2000). They recommended increasing observer coverage to 56-83% in order to obtain a robust estimate of bycatch (CV of 30%; Baird and Bradford 2000). Unfortunately, more recent observer programmes have been highly localised and typically had very low levels of observer coverage, typically less than 5% (Tables 3-5).

**Table 3.** Observer coverage in the New Zealand inshore gillnet fishery (Abraham and Thompson 2011; Rowe 2009, 2010; Ramm 2010, 2012; Clemens-Seely et al. 2014a, b; Clemens-Seely and Hjorvarsdottir 2016).

| Fisheries Management Area | Observer coverage |  |  |  |  |  | 2012-13 | 2013-14 | 2014-15 |
| --- | --- | --- | --- | --- | --- | --- | --- | --- | --- |
|  | 1999-00 | 2000-01 | 2005-06 | 2007-08 | 2008-09 | 2010-11 |  |  |  |
| 1 |  |  |  |  |  |  |  |  |  |
| 2 |  |  |  |  |  | 0.006 |  |  | 0.002 |
| 3 |  |  | 0.0043 | 0.068 | 0.215 | 0.098 | 0.019 | 0.016 | 0.021 |
| 4 |  |  |  |  |  |  |  |  |  |
| 5 |  |  | 0.052 | 0.250 | 0.240 |  |  |  | 0.368 |
| 6 |  |  |  |  |  |  |  |  |  |
| 7 |  |  | 0.051 | 0.001 | 0.070 |  |  | 0.011 |  |
| 8 |  |  | 0.002 | 0.048 |  |  | 0.278 | 0.352 | 0.267 |
| 9 |  |  |  | 0.002 |  |  | 0.001 |  | 0.001 |
| Nationwide | 0.001 | 0.001 | 0.004 | 0.025 | 0.043 | 0.019 | 0.022 | 0.024 | 0.034 |

Two estimates of Maui dolphin bycatch are available, based on the remaining overlap of gillnet and trawl fisheries with Maui dolphin distribution. A government convened Expert Panel of national and international scientists estimated an annual bycatch of 4.97 Maui dolphins (95% CI 0.28-8.04), 3.83 in gillnets and 1.13 in trawl fisheries (Currey et al. 2012). Slooten (2014) updated this estimate after the area where gillnets are banned was extended in 2012 and 2013, and estimated current bycatch at 3.25-4.16 Maui dolphins per year (1.13 in trawl fisheries and 2.15-3.03 in gillnets). Observer coverage in Maui dolphin habitat off the WCNI is 14.6% for trawling vessels (IWC 2016). Observer coverage for gillnetting vessels in Maui dolphin habitat is estimated at 12.7 % for vessels > 6 m (IWC 2016) but only 2% when vessels < 6m in length are included (MPI 2016b).

### PBR estimates

Updated estimates of PBR for regional populations of Hector’s dolphins are shown in Table 6. We applied the standard approach of using N_min_ estimates for populations which are small enough to ensure that there is only one population and one fishery in each region. We used a recovery factor of 0.1 given that Hector’s dolphin is listed as Endangered and Maui dolphin as Critically Endangered by both national (Department of Conservation) and international agencies (IUCN).

**Table 4.** Observer coverage in the New Zealand inshore trawl fishery (Rowe 2009, 2010; Ramm 2010, 2012; Clemens-Seely et al. 2014a, b; Clemens-Seely and Hjorvarsdottir 2016). Notes: *15 trawls observed out of a total of 15 trawls.

| Fisheries Management Area | Observer coverage |  |  |  |  |  |  |  |  |
| --- | --- | --- | --- | --- | --- | --- | --- | --- | --- |
|  | 2006-07 | 2007-08 | 2008-09 | 2009-10 | 2010-11 | 2011-12 | 2012-13 | 2013-14 | 2014-15 |
| 1 | 0.009 |  | 0.026 | 0.017 | 0.013 | 0.007 | 0.004 | 0.121 | 0.171 |
| 2 | 0.001 |  |  |  | 0.036 |  |  | 0.008 | 0.014 |
| 3 |  | 0.004 | 0.071 | 0.028 |  | 0.021 |  | 0.018 |  |
| 4 |  |  |  |  |  |  | 1.00* |  | 0.003 |
| 5 | 0.001 |  | 0.048 | 0.051 | 0.002 |  |  |  |  |
| 6 |  |  |  |  |  |  |  |  |  |
| 7 | 0.001 | 0.005 | 0.039 | 0.016 | 0.017 | 0.050 |  | 0.012 |  |
| 8 |  | 0.005 | 0.020 |  | 0.006 |  | 0.001 | 0.016 | 0.013 |
| 9 | 0.017 | 0.018 | 0.039 |  |  | 0.042 | 0.002 | 0.079 | 0.198 |
| Nationwide | 0.003 | 0.003 | 0.035 | 0.018 | 0.013 | 0.025 | 0.001 | 0.032 | 0.041 |

**Table 5.** Gillnet fishing effort, proportion of effort observed and observed Hector’s dolphin captures from Abraham et al. (2016).

| Fishing year | Fishing effort | Observed effort | Observed % | Observed captures |
| --- | --- | --- | --- | --- |
| 2002/2003 | 28,268,690 |  |  |  |
| 2003/2004 | 27,052,566 |  |  |  |
| 2004/2005 | 27,199,843 |  |  |  |
| 2005/2006 | 24,257,075 | 127,778 | 0.005 | 0 |
| 2006/2007 | 24,268,896 | 167,060 | 0.007 | 1 |
| 2007/2008 | 21,198,516 | 295,400 | 0.014 | 1 |
| 2008/2009 | 20,942,804 | 501,405 | 0.024 | 1 |
| 2009/2010 | 22,258,289 | 581,689 | 0.026 | 2 |
| 2010/2011 | 21,663,381 | 174,500 | 0.008 | 0 |
| 2011/2012 | 19,965,213 | 86,892 | 0.004 | 0 |
| 2012/2013 | 21,705,826 | 732,070 | 0.034 | 1 |
| 2013/2014 | 20,889,916 | 214,100 | 0.010 | 1 |

**Table 6.**
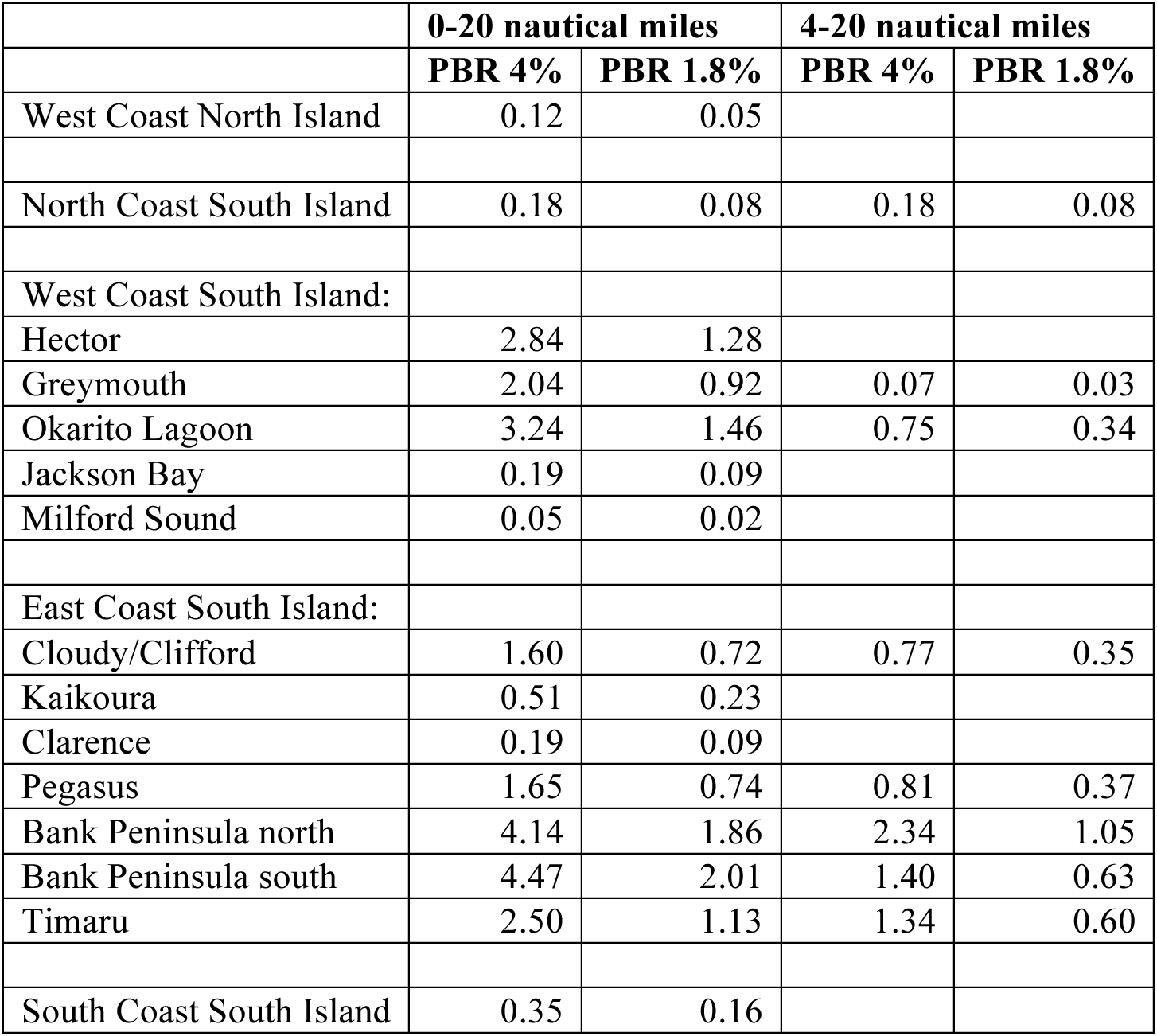
Potential Biological Removal (PBR) for regional populations of Maui (west coast North Island) and Hector’s dolphins (all other populations), estimated following the methods of Wade (1998). We used two offshore ranges: Shoreline to 20 nautical miles offshore (full offshore range for which population estimates are available) and 4-20 nautical miles (proportion of the population outside protected areas) and two levels of Rmax, the default value of 4% for cetaceans and the 1.8% Rmax estimate for Hector’s and Maui dolphins.

Table 6 provides PBR estimates for the default Rmax of 4% for cetaceans as well as the Rmax of 1.8% estimated for Hector’s and Maui dolphins. We have provided estimates for the total population in regional areas as well as the population outside protected areas. Note: If animals inside protected areas were to be included in the PBR calculation, this would undermine the effectiveness of the protected areas. PBRs for most populations are less than one individual per year, with a minimum of 0.03 – 0.07 for the offshore Greymouth region on the WCSI and a maximum of 2.01 – 4.47 dolphins per year for the full distribution off southern Banks Peninsula on the ECSI. These PBR estimates compare with an estimated 23 dolphins caught per year caught in gillnet fisheries off the east coast of the South Island (Slooten and Davies 2011). PBRs for the whole South Island range from 3 to 24, depending on the value used for R_max_, and which offshore range is used. The total PBR for the ECSI is 3 – 15. Recent estimates of 110 – 150 Hector’s and Maui dolphins caught per year during the years 2000-2006 were clearly far in excess of sustainable levels (Davies et al. 2008).

### Results of population viability and power analysis

The new Population Viability Analysis (PVA) indicates that the 1975 population size of South Island Hector’s dolphins was on the order of fifty thousand individuals (50,158 95%, CI 27,411 – 91,783, Table 7). Current population size is estimated at 30% of the 1975 level. This equates to a 70% decline over the last three generations (39 years), compared to the previous estimate of a 74% decline (IUCN 2008). The IUCN criterion for Endangered is a population decline > 50% over three generations.

**Table 7.**
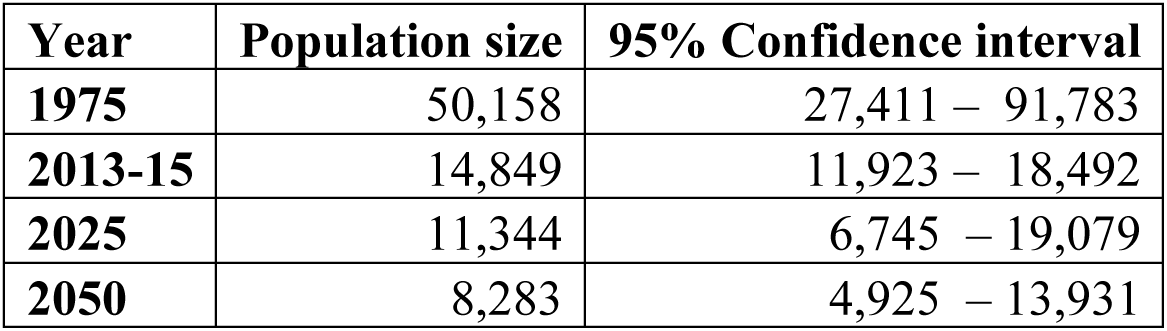
The most recent population estimate for South Island Hector’s dolphins compared with estimates from the population viability analysis for past (1975) and future population sizes (2025 and 2050).

| Year | Population size | 95% Confidence interval |
| --- | --- | --- |
| 1975 | 50,158 | 27,411 – 91,783 |
| 2013-15 | 14,849 | 11,923 – 18,492 |
| 2025 | 11,344 | 6,745 – 19,079 |
| 2050 | 8,283 | 4,925 – 13,931 |

Comparing these new results with previous PVAs (Slooten and Dawson 2010) indicates that the high level of overlap between gillnet (and trawl) fisheries and the range of Hector’s dolphin more than offsets the apparently larger population size off the ECSI. The prognosis for Hector’s dolphin remains essentially the same. The results of this updated PVA are similar to scenarios from Slooten and Dawson (2010) that included displacement of fishing effort from protected areas to areas that still have relatively high densities of Hector’s dolphins. This makes sense in the light of the most recent information on offshore distribution of Hector’s dolphins. Comparing population estimates for the area from 0-4 nautical miles offshore (the area covered by the previous surveys) suggests a population decline of c. 1000 individuals for WCSI. This matches the population decline predicted in earlier population viability analyses (Slooten and Dawson 2010).

The linear regression of the natural log of the abundance estimates for Maui dolphin, followed by back-transformation, indicated a decline from about 138 Maui dolphins in 1985 to 56 individuals in 2016 (Figure 7). This is an annual decline of 2% and a total decline of 59% over the 31 year period from 1985 to 2016. The 95% confidence interval of the difference between the last two Maui dolphin population estimates range from a 1.6% decline per year to a 4.8% increase per year. This indicates a high level of uncertainty about whether the population is increasing, decreasing or stable. The Bayesian linear regression of the last three estimates indicated a 68% probability of population decline.

**Figure 7.**
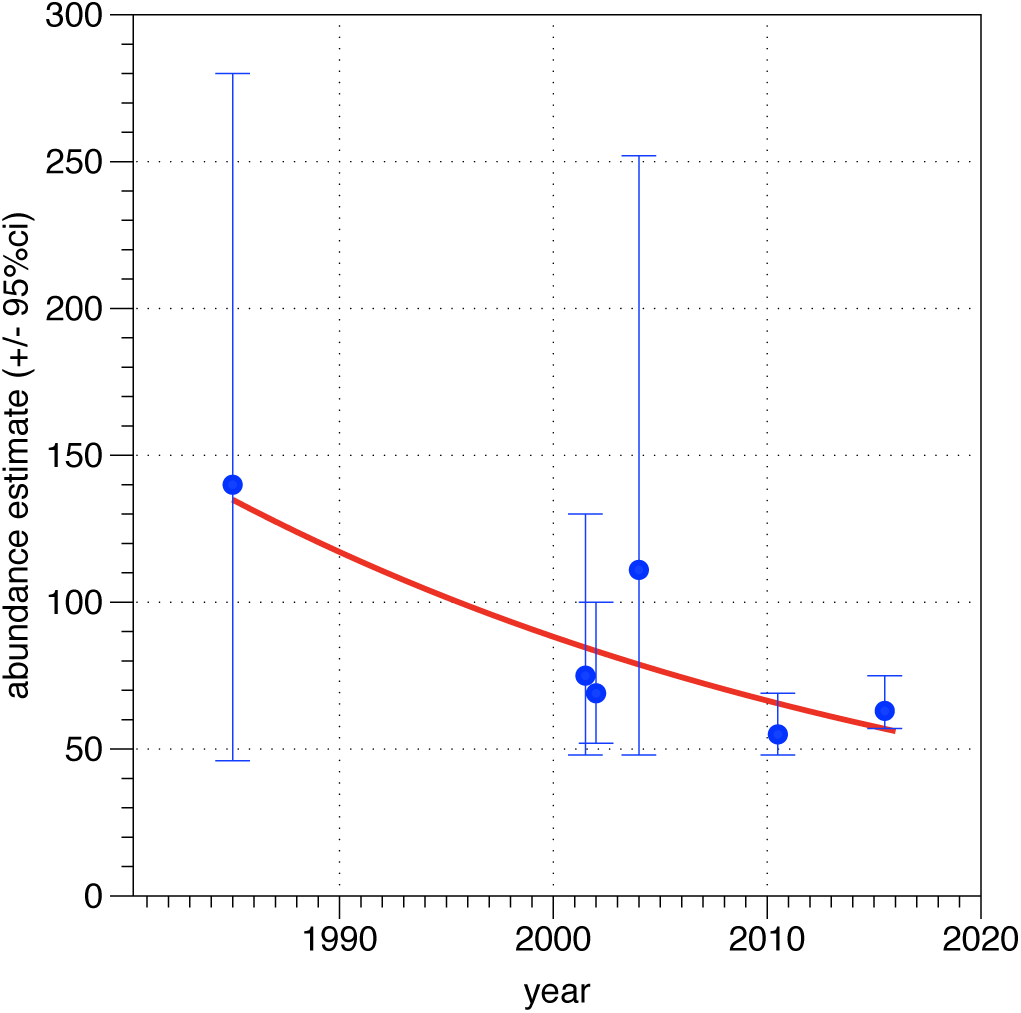
Linear regression of log-transformed abundance estimates, back-transformed to original scale. Abundance estimates are from (Dawson and Slooten 1988, Ferreira and Roberts 2003, Slooten et al. 2006, Hamner et al. 2012, Baker et al. 2012, 2016).

We estimated the statistical power of being able to detect a population recovery of 1.8% per year or a continued population decline of 3% per year (the rate of decline estimated by Wade et al. 2012, which will be easier to detect than our estimated decline of 2% per year). The results indicate that statistical power for detecting Maui dolphin population trends is very low (Figure 8, Table 8). After 30 years (6 more surveys) the statistical power of determining if the population is continuing to decline at 3% per year would be 13% and the power of detecting a realistic rate of population recovery (1.8% recovery per year) would be 7%. Even with multiple surveys, only fairly large population changes will be detectable. Thirty years of surveys every five years will be required to detect with 80% statistical power a population increase of 45%, or decline of 37%.

**Table 8:** Estimates of statistical power, starting with a population of 63 Maui dolphins (CV 0.09) with surveys conducted every five years, as at present. The last two columns show the level of population icrease and decrease (effect size) detectable with 80% power.

| Number of surveys | Number of years | Power of detecting 1.8% per year increase | Power of detecting 3% per year decrease | Decline over 5 years detectable with 80% power | Recovery over 5 years detectable with 80% power |
| --- | --- | --- | --- | --- | --- |
| 3 | 15 | 0.05 | 0.06 | >1 | >1 |
| 4 | 20 | 0.06 | 0.09 | 0.59 | 0.81 |
| 5 | 25 | 0.06 | 0.11 | 0.43 | 0.55 |
| 6 | 30 | 0.07 | 0.13 | 0.37 | 0.45 |

**Figure 8:**
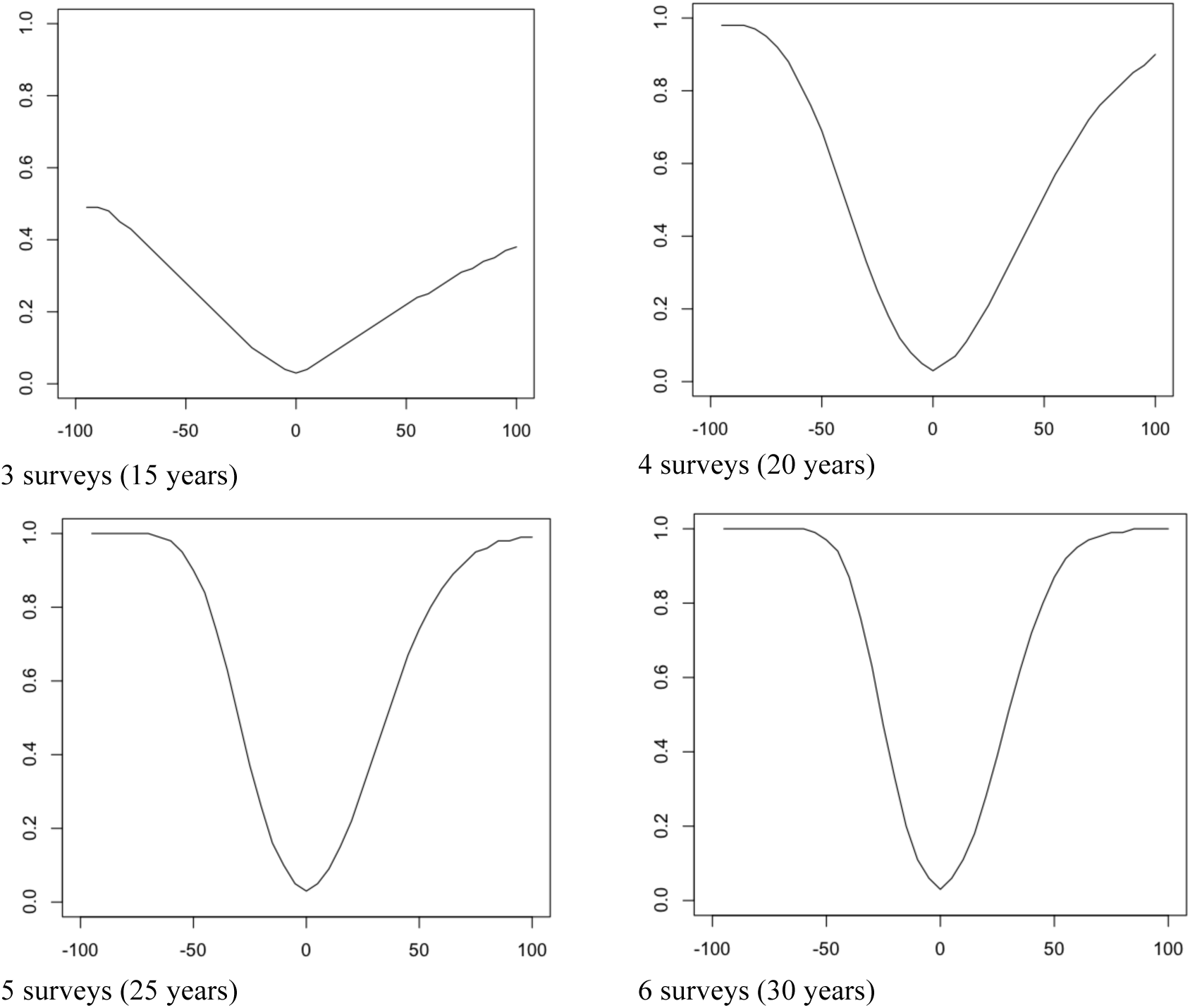
Statistical power (y axis) to detect a given population trend (x axis) after 3, 4, 5 and 6 surveys (one survey every 5 years).

## DISCUSSION

Bycatch estimates for the commercial gillnet fishery are available for a small number of years and regions, starting with an observer programme conducted at Banks Peninsula during the 1997-1998 fishing season (Baird and Bradford (2000). No estimate is available for the number of Hector’s or Maui dolphins caught in trawl fisheries or recreational gillnet fisheries. They have been observed caught in trawl nets (e.g. Baird and Bradford 2000) and recreational or amateur gillnets (DOC and Mfish 2007), but observer coverage has been too low to allow estimation of the catch rate. Therefore the analyses in this paper are based on commercial gillnet bycatch only, and are inevitably optimistic.

Baird and Bradford (2000) recommended an increase in the level of observer coverage in the gillnet fishery to > 55% in order to achieve a CV on the bycatch estimate of 30% or better. Subsequent observer programmes have not been designed to achieve a specific level of precision or accuracy, and have had very low coverage. Scientists involved with estimating fisheries mortality have repeatedly pointed out that observer coverage is too low to allow estimation of Maui and Hector’s dolphin bycatch (e.g. Baird and Bradford 2000; Dawson and Slooten 2005; Rowe 2009; Abraham et al. 2010; Thompson et al. 2013).

Bycatch in trawl fisheries poses an even greater challenge, because the strike rate per trawl is likely to be very low. However, the very large number of trawls means that this source of mortality could be considerable and cannot be ignored (Dawson and Slooten 2005). According to Rowe (2009) “The extent to which inshore trawl vessels interact with protected species is extremely poorly known due to minimal historic observer coverage in almost all areas.”

Davies et al. (2008) used the Baird and Bradford (2000) data to carry out a more detailed analysis, which took account of water depth, target fish species and other variables. They estimated a total bycatch level during 2000-2006 of 110 to 150 Hector’s and Maui dolphins per year. Davies et al. (2008) estimated that 35-46 Hector’s dolphins per year per year were caught off the east coast of the South Island. This is the only area for which two estimates are available, from before and after 2008 when protection from gillnet and trawl fisheries was extended. Slooten and Davies (2011) estimated that the extended protection measures implemented in 2008 resulted in a reduction of Hector’s dolphin bycatch off the east coast of the South Island from 35-46 per year before 2008 to 23 per year (CV 0.21) after 2008 and a corresponding reduction in the rate of population decline. No such estimates are available for other parts of New Zealand waters. The relatively high level of continuing bycatch off ECSI is consistent with independent data on Hector’s dolphin survival rates from Banks Peninsula (Gormley et al. 2012). Research using photographic capture-recapture analysis shows a 5.4% improvement in survival rates in this area – one of the ECSI hotspots for Hector’s dolphins. Partial protection of this population has reduced a 6% per year population decline to a decline of around 1% (Gormley et al. 2012). The probability of population recovery was recently estimated at 41%, compared to 7% previously (Gormley et al. 2012).

Observer programmes, with sufficiently high coverage to result in scientifically robust estimates of bycatch, are urgently needed in at least two additional areas. Expanding geographic coverage would put calculation of total catches on a much more robust footing. The obvious areas for these observer programmes are the west and south coasts of the South Island. These areas have relatively high dolphin densities and therefore provide a high probability of obtaining statistically robust data, providing that observer coverage levels are reasonable. Maui dolphin densities off the west coast of the North Island are extremely low, with a population of fewer than 70 individuals spread over a very large area. An observer programme in this area would waste resources and distract attention from the obvious necessity that, for this population to persist, let alone recover, bycatch must be essentially eliminated.

MPI is currently planning to increase the use of video camera monitoring on fishing vessels to improve the accuracy of fish catch and bycatch reporting. Such systems hold considerable promise to increase the “observer” coverage of NZ’s inshore fleet. However, the effectiveness of video monitoring is dependent on the “view” and image quality of the system, the system’s reliability, the skill and diligence of the people employed to view the resulting video records. Appropriate follow-through by the authorities is also required, and has been lacking (e.g. MPI 2013). For the data to be credible, it is essential that the analysis of the video records is independent of the fishing industry (currently a fishing industry owned company carries out video monitoring).

Even if on-board video systems meet all expectations and are employed on all fishing vessels, there will be a need for on board observers to estimate drop-out (e.g. Tregenza et al. 1997; Hamer et al. 2011; Uhlmann and Broadhurst 2015). A recent video camera trial showed a fisherman letting out a gillnet in an effort to promote drop out. One dolphin was retained in the net, and was reported. They other dropped out of the net (after the net was left in the water for almost an hour) and was not reported.

Other problems identified in recent video camera trials in New Zealand include fishermen turning the cameras off (McElderry 2007; MPI 2013), and camera failure. In a current trial of video observation in New Zealand’s snapper fishery, 80% of the cameras are reported to have failed. Most of these problems can be solved. Camera systems can be set up to activate automatically when the net is hauled and designed so they cannot be manually switched off. Cameras with a wide field of view may help ensure drop-out can be documented but will offer less detail in any one part of the image, perhaps compromising species identification or condition assessment.

By far the largest problem is the time taken to review footage, and the question of who does that task. Restricting oversight to a fraction of the footage, as is done in New Zealand, raises the possibility of bias. For example, MPI (2013) automatically rejected hauls in excess of 3 hours, due to the length of time it would take to view them. Review of footage gathered in upcoming camera deployment has been contracted to a company wholly owned by the fishing industry.

Abraham and Thompson (2011) concluded that “reporting of captures by fishers was at a lower rate than reporting of captures by observers”. This is also true in the Auckland Islands trawl fishery for squid, which has a significant bycatch of NZ sealions. Research on fish bycatch also shows a strong observer effect, with fishing vessels behaving very differently depending on whether they were carrying observers or not (Bremner et al. 2009; Simmons et al. 2016). A key feature of the way in which Marine Mammal bycatch is quantified in New Zealand is the existence of several biases which together act to substantially underestimate the true numbers taken.

The results of the updated PVA are consistent with previous risk analyses. The current populations of Hector’s and Maui dolphins are on the order of 30% and 10%, respectively of their population size in 1975. At 70% the rate of decline for Hector’s dolphin over three generations exceeds the IUCN threshold for Endangered and is very similar to the previous estimate of the rate of decline (74%, IUCN 2008).

Population projections to 2025 and 2050 predict continued population declines under the current protection measures. All risk analyses to date show that without fisheries mortality, Hector’s and Maui dolphins are predicted to recover, potentially up to half of their original population size (OSP) by 2050 (Slooten and Dawson 2010, Slooten and Davies 2011). Our PVA based on the most recent population estimate indicates that the increased overlap between Hector’s dolphins and fishing methods known to cause dolphin bycatch (gillnets and trawling) outweighs the apparent increase in population size.

We agree that the extremely small population of Maui dolphin qualifies the sub-species for Endangered status under the ESA. The 68% probability of continued population decline for Maui dolphins is not encouraging. Continued population monitoring has very low statistical power to detect population recovery or continued declines. This means that for the foreseeable future a precautionary approach will be necessary. The IWC has recommended switching to fishing methods that do not cause dolphin mortality throughout the full range of Maui dolphins (IWC 2016). The IUCN has recommended banning gillnet and trawl fisheries throughout the range of Hector’s and Maui dolphins (IUCN 2016).

The genetic mark-recapture surveys should not be thought of as representing the whole Maui dolphin population. This is for two reasons. First, the Department of Conservation does not allow biopsy of juvenile dolphins, thought to be less than 1 year old. Therefore, the mark-recapture estimates do not include this part of the population. Additionally, almost all biopsy sampling effort has occurred in the centre of the current Maui’s dolphin distribution, near port Waikato. While attempting to biopsy exceptionally rare animals near the limits of their range is likely to be frustratingly unproductive, dolphins near those limits are unlikely to have been sampled. This makes it very difficult to detect reductions in the range of Maui dolphins over time.

### Concluding remarks

The reported bycatch of NZ dolphin substantially understates the actual level of bycatch, due to very low levels of self reporting by fishermen and low observer coverage. To date, almost all bycatch information has come from observer programmes off the east coast of the South Island. Observer programmes off the west and south coasts of the South Island would help determine if bycatch rates (per unit of fishing effort and dolphin density) differ significantly from one area to another. Scientifically robust estimates of bycatch for two additional areas would substantially improve the state of knowledge about Hector’s and Maui dolphin bycatch. Observer coverage off the west coast of the North Island has no realistic chance of resulting in an estimate of bycatch. Likewise, the very low statistical power for detecting Maui dolphin population trends makes it impractical to monitor the population in the hope of determining whether the current, partial protection is effectiveness. The focus for Maui dolphins should be to improve protection, rather than gathering more research data. The IWC Scientific Committee has recommended such a precautionary approach since 2012. The results of the population viability analysis and PBR calculations are consistent with earlier analyses. These updated analyses confirm the IUCN listing of Hector’s dolphin as Endangered an Maui dolphin as Critically Endangered. These results support the IUCN recommendation to avoid the use of gillnets and trawling throughout the range of Hector’s and Maui dolphin and NOAA’s proposal to list these taxa under the Endangered Species Act.

## Acknowledgements

We would like to thank Lindsay Wickman for help with the Bayesian linear regression.

